# Quantification of Hypoperfusion in Autism Assessed by Diffuse Correlation Spectroscopy and Frequency Domain Near Infrared Spectroscopy

**DOI:** 10.1101/2020.11.05.369918

**Authors:** Benjamin Rinehart, Chien-Sing Poon, Ulas Sunar

## Abstract

There is a need for quantitative biomarkers for early diagnosis of autism. We showed previously that hemoglobin concentrations show contrast between autistic group vs control group in humans. Cerebral blood flow can provide additional sensitivity for improved characterization. Diffuse correlation spectroscopy (DCS) has been shown to be reliable method to obtain blood flow contrast in animals and humans. Thus, in this study we evaluated DCS in an established autism model. Our results indicate that autistic group had significantly lower blood flow compared to control group. These results confirm the previous results obtained from humans by positron emission tomography and magnetic resonance imaging.

## 1. Introduction

Autism spectrum disorders (ASD) is now recognized to occur in approximately 1% of the population and is a major public health concern because of their early onset, lifelong persistence, and high levels of associated impairment [1]. ASD is a highly heritable, biologically based neurodevelopmental disorder. Despite this fact, the exact cause remains unknown. Finding the cause has been challenging because ASD encompasses a range of complex disorders that involve multiple genes and demonstrate great phenotypical variation. Estimates of recurrence risks based on family studies of idiopathic ASD are approximately 6% when there is an older sibling diagnosed with ASD, and even higher when there are already two children with ASDs in the family [2]. One of the main difficulties when diagnosing and treating autism is the observational nature of the diagnostic schedules [3]. Although these diagnostic methods have a high sensitivity of around 90%, the specificity can be quite low, leading to overdiagnosis and overtreatment. Also, the time requirement is cost-intensive and requires a dedicated team of physicians to accurately track progress.

There exists a need for a non-observational diagnostic system capable of measuring in-vivo biomarkers to both accurately place a patient along the ASD spectrum and track the efficacy of therapies in patients that receive them. Recent in-vivo imaging studies have shown that cerebral hypoperfusion presents as a comorbidity in ASD subjects when compared with control groups. Traditionally these measurements have been performed using positron emission tomography (PET) or single-photon emission computed tomography (SPECT) [4–8]. Functional magnetic resonance imaging (fMRI) was also used in a series of ASD neuro-imaging studies with a sensitivity of ASD diagnosis up to 75% [9–11]. An arterial spin labeling MRI study also showed robust hypoperfusion results in the ASD population [12]. The results of these studies are encouraging, however, the imaging modalities of PET, SPECT, and MRI are limited for routine clinical use in adolescent ASD diagnosis. The extended imaging durations and the sensitivity to motion artifact also present a risk in rendering data unusable.

Efforts have been made in the optical imaging field to fill this need as optical imaging techniques possess the unique combination of advantages of being relatively inexpensive, simple to operate, and with non-contact imaging options. Initially, functional near-infrared spectroscopy (fNIRS) has been used to monitor hemodynamic abnormalities in autism subjects through monitoring changes in blood chromophore concentrations (specifically oxygenated and deoxygenated hemoglobin) as we have previously shown in clinical studies [13,14]. Laser Doppler flowmetry was performed as part of a multimodal imaging study which found neurovascular dysfunction in the 16p11.2 deletion mouse model of ASD [15]. Laser speckle imaging (LSI) at 532nm was applied for an autism model in mice with the advantage of wide-field imaging in a non-contact, camera-based setup [16]. However, it has limited penetration depths of 100s of microns, where majority of the reflected signal will be from the scalp tissue. Therefore, there exists a need for a low-cost, non-invasive optical approach that is capable of measuring brain blood flow in the ASD mouse model. Here, we present results from a custom-built diffuse correlation spectroscopy (DCS) system that we used previously in our human studies [17].

## 2. Material and methods

### 2.1 Animals

Three-month-old mice were sourced from Jackson Labs (Bar Harbor, ME, USA). Study groups contained 6 male C57BL/6J (control) and 6 age-matched male BTBR T+ Itpr3tf/J (ASD) mice. The groups will hereby be referred to using their common names of C57/B and BTBR, respectively. The BTBR mouse model is previously phenotyped to show similar symptoms and behaviors to the human form of ASD [18,19]. It is believed that the structural abnormality of an absent corpus callosum is a key component associated with core behavioral issues of ASD [20,21].

The heads of the mice were shaved, and a chemical depilatory agent was applied to ensure that the imaging area was clear of hair. The hair removal procedure was done 24 hours in advance of imaging to allow time for any redness or swelling caused by minor skin irritation to subside.

Immediately prior to imaging, the mice were anesthetized through gaseous administration of an isoflurane and oxygen blend. To control for the decreased blood flow caused by hypertension as a result of isoflurane administration all of the mice were anesthetized and imaged on a precise schedule [22]. Each mouse was incubated with 5% isoflurane at a flow rate of 300 ml/min for 2 minutes. The mice were then transferred to a nose cone where the isoflurane dose was dropped to 2.5% at the same flow rate. DCS measurements were taken starting at 7 minutes after initial anesthesia administration and frequency domain measurements followed starting at 9 minutes.

All procedures related to animal care were conducted with the approval of the Institutional Animal Care and Use Committee at Wright State University.

### 2.2 Diffuse Correlation Spectroscopy

In-depth details of the DCS technique have been covered in previous reviews [23–25]. The DCS instrument used in this study has been described in our previous work [16,25]. Briefly, DCS was initially used for monitoring flow dynamics in a turbid medium, such as living tissue [24,27–30]. DCS measures the emitted light intensity autocorrelation function, whose decay rate reflects the dynamics of “scatterers” - in this case moving blood cells in tissue [23,27]. The dynamic information of the scatterers (moving blood cells in the microvasculature) can be obtained by fitting the model to the experimental data [22,23,26]. It was shown that the diffusive motion, αD_B_, can model the dynamics in deep tissue to obtain the blood flow index (BFI), where α is proportional to tissue blood volume fraction, and D_B_ is the effective Brownian diffusion coefficient [26,27,31]. The normalized diffuse electric field temporal autocorrelation function (g1 (r,τ)) extracted from normalized intensity temporal autocorrelation function (g2 (r,τ)) via Siegert relation is fitted to an analytical solution of the diffusion equation to estimate the BFI parameter.

In this study, the DCS system consists of a continuous-wave laser source (785 nm CrystaLaser, Reno, NV, USA) with coherence length longer than 10 m, eight NIR-optimized single-photon counting module (SPCM-NIR, Excelitas, QC, Canada), and a 8-channel auto-correlator board (Correlator.com), of which one channel was used. A single multi-mode fiber (1000 um core diameter, 0.39 numerical aperture (NA)) was used to guide the 785 nm laser light to the scalp and a single-mode fiber (5 um core diameter, NA of 0.13) collected the light emitted from the scalp to the single-photon counting modules, as shown below in (Fig 1A). The separation distance of the source and detector fiber was set at 5 mm with the detector fiber placed at the midpoint of the cerebral cortex and a translational stage was used to place the source fiber 5 mm anterior to the detector fiber, as shown below (Fig 1B).

**Figure 1.**
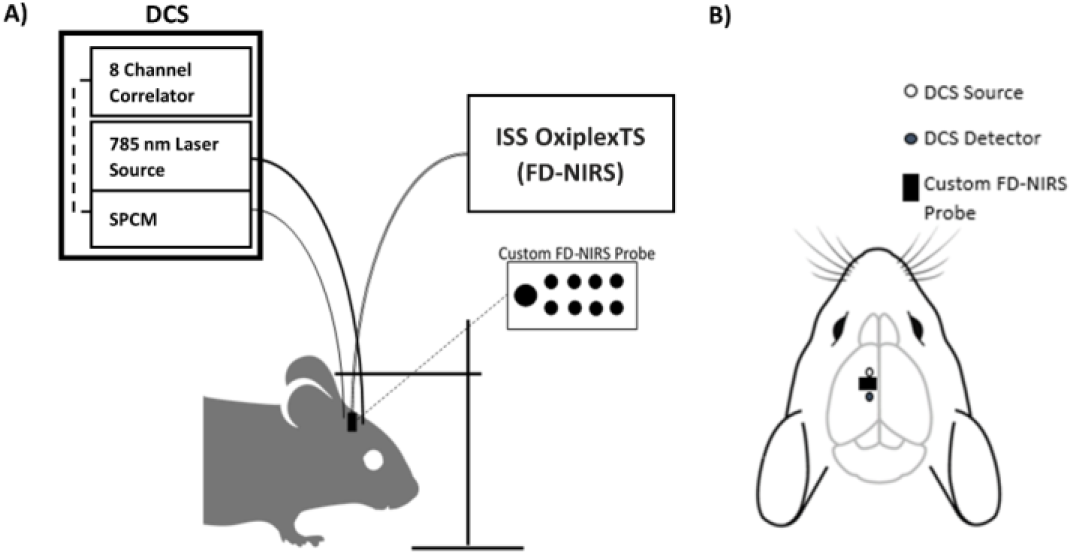
(A) Optical instrumentation of DCS and FD-NIRs systems guiding optical information to and from the mouse scalp. (B) Top down view of the probe placement on the scalp in reference to the mouse brain underlying the scalp.

The optical parameters (absorption coefficient, µ_a_, and reduced scattering coefficient, µ_s_’) at 690 nm and 830 nm were resolved using a frequency-domain NIRS system (OxiplexTS, ISS Inc., Champaign, IL, USA). A custom probe with source-detector separations at 2, 4, 6, and 8 mm was fabricated using 400 um multimode source fibers and a single 1000 um detector fiber to allow probing optical property measurements at a similar depth as our DCS measurements. The optical properties measured at 830 nm were used as inputs for the BFI calculations from the DCS data.

### 2.3 Immunohistochemistry

Previous reports have shown that the neurogenesis markers of doublecortin (DCX) and Ki-67 are positively associated with CBF decreases in the BTBR mouse model measured with DW-LSI [13]. DCX is a microtubule-associated protein expressed by neuronal precursor cells and immature neurons in adult cortical structures and Ki-67 is detectable within the nucleus during proliferation. Therefore, we measured against this same biomarker using the immunohistochemistry (IHC) protocol outlined via Abookasis et al. [16] with primary and secondary antibodies sourced from Abcam (Cambridge, United Kingdom) DCX primary 18723, Ki-67 primary 15580. The DCX sections were stained with a primary concentration of 1:1000 in 0.5% bovine serum and a secondary concentration of 1:250. The Ki-67 sections were stained with a primary concentration of 1:200 in 0.5% bovine serum and a secondary concentration of 1:250. Stained sections were imaged under compound light microscopy at x400 magnification. DCX sections were imaged in the granular cell layer (GCL) of the dentate gyrus (DG) and the Ki-67 sections were imaged in the sub-granular zone.

## 3. Results and discussion

Optical absorption coefficient (µ_a_) and reduced scattering coefficient (µ_s_’) results at 690 nm and 830 nm are summarized for BTTR and C57/B6 groups in Figure 2. For µ_a_ measurements (Fig 2A), the BTBR group had 0.244 ± 0.007 at 690nm, and 0.224 ± 0.013 cm^−1^ at 830 nm, while these values were 0.223 ± 0.003 cm^−1^ and 0.203 ± 0.008 cm^−1^ for the C57/B6 group. The Wilcoxon rank sum test showed that the differences in µ_a_ between the groups were significant, p = 0.0036 at 690 nm and p = 0.0246 at 830 nm. Reduced scattering coefficient (µ_s_’) measurements for BTBR 690 nm was 8.38 ± 0.20 cm^−1^, and 7.00 ± 0.90 cm^−1^ at 830 nm, and for C57/B6 µ_s_’ at 690 was 8.91 ± 0.33 cm^−1^ and 6.42 ± 0.30 cm^−1^ showed that 830 nm. The Wilcoxon rank sum test showed that the difference was not statistically significant (p>0.05).

**Figure 2.**
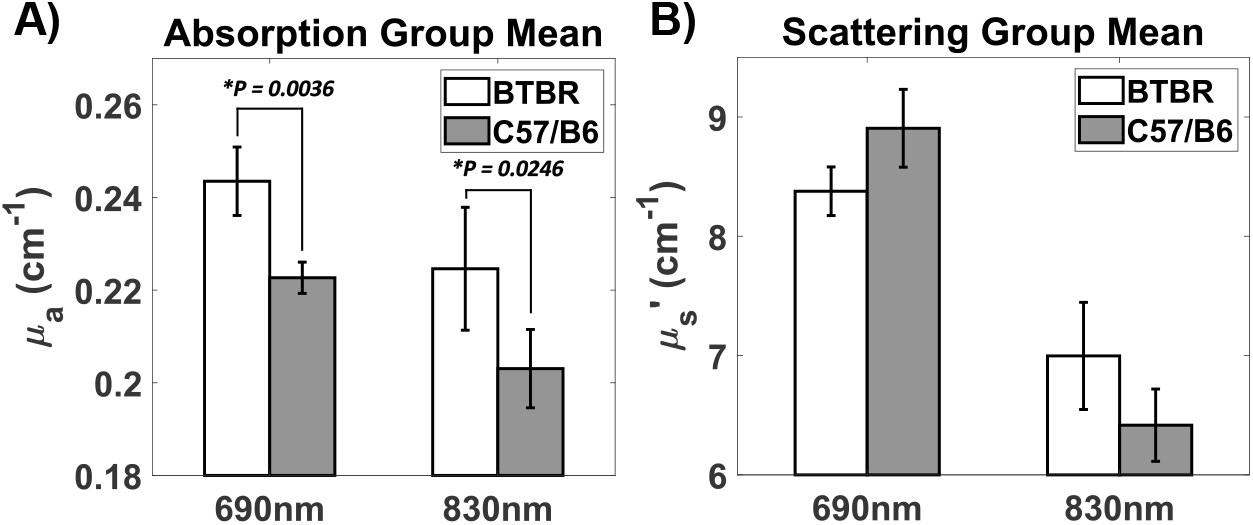
(A) Absorption coefficient (µ_a_) measurements for BTBR (690 nm, 830 nm) and C57/B6, indicating significantly lower absorption at 690 nm (p = 0.0036) and 830 nm (p = 0.0246) for the C57/B6 group. (B) Reduced scattering coefficient (µ_s_’) measurements for BTBR (690 nm, 830 nm) and C57/B6.

The individual blood flow index results in Figure 3A and the group mean values are shown in Figure 3B. The group average of αD_B_ measurements for BTBR was (0.56 ± 0.02) * 10^−8^ cm^2^/s, while it was (1.12 ± 0.06) * 10^−8^ cm^2^/s for C57/B6 mice. The Wilcoxon rank sum test showed that there was a significant difference (p = 0.0152) between the BTBR and C57/B groups, clearly indicating that there exist hypoperfusion in the BTBR group compared to the control group.

**Figure 3.**
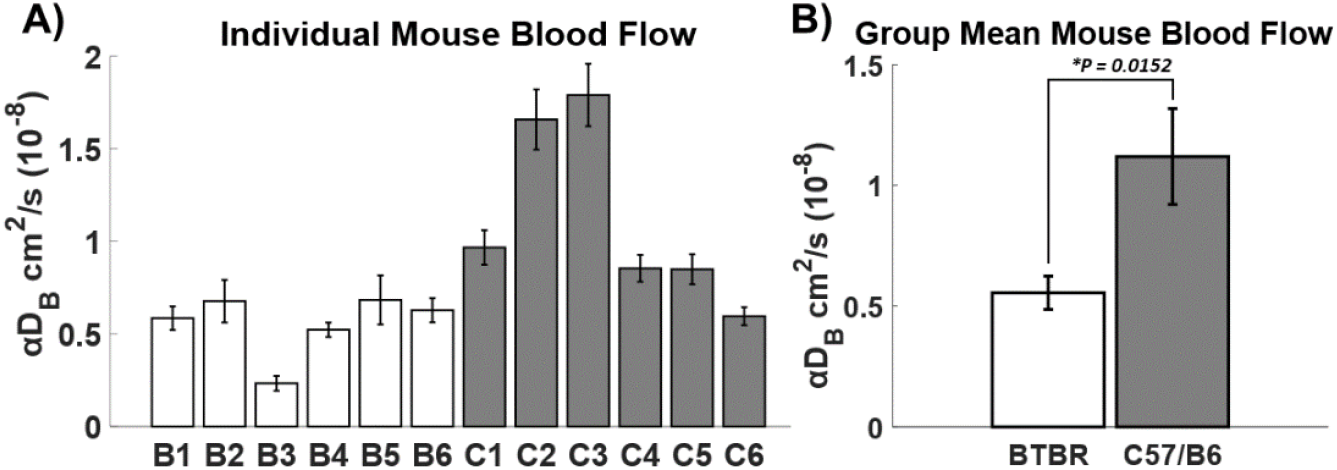
(A) Individual blood flow measurements of each mouse in terms of (αD_B_). (B) Group average of αD_B_ measurements for BTBR (0.56 +/- 0.02) * 10^−8^ cm^2^/s and C57/B6 (1.12 +/- 0.06) * 10^−8^ cm^2^/s. A significantly lower blood flow (hypoperfusion) is observed for the BTBR group (p = 0.0152).

The results of the IHC staining on the brain sections are shown below (Fig 4). The DCX images show a higher cell count in the representative control image (Fig 4A) as compared to the representative autism tissue image (Fig 4C). The Ki-67 staining of the autism image appears to have an underdeveloped sub-granular zone (Fig 4D), causing the cell count in the control sections (Fig 4B) to be higher by default.

**Figure 4.**
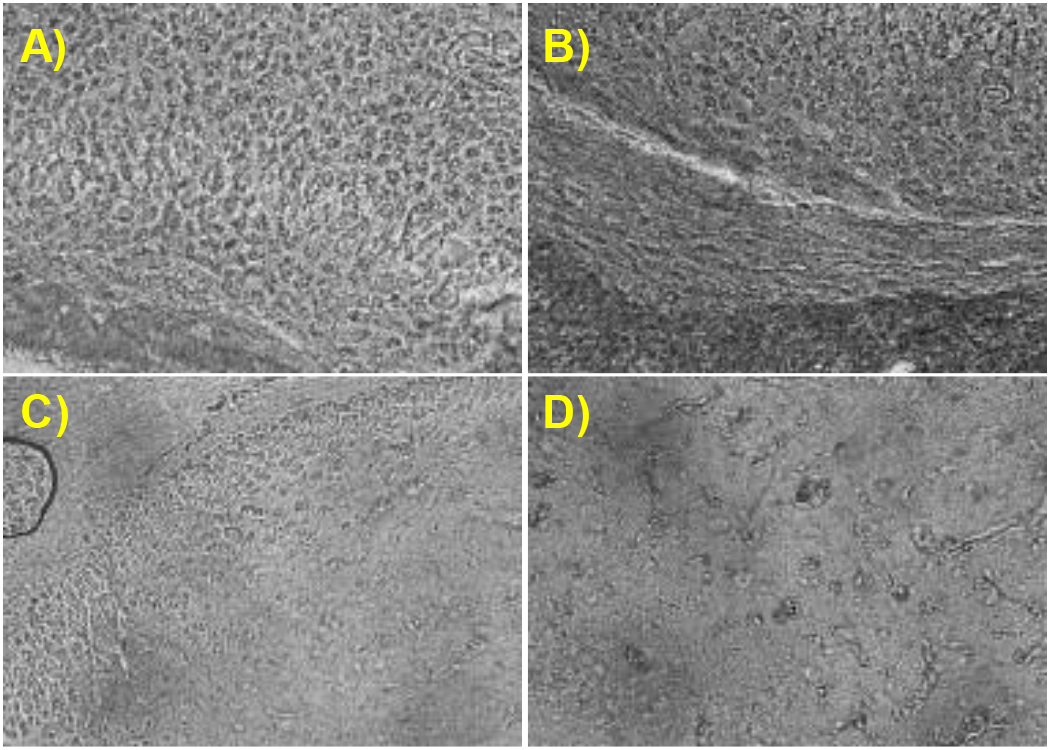
DCX IHC staining results of the (A) C57/B6 and (C) BTBR representative sections. Images were centered on the GCL of the DG at 400x magnification. Ki-67 IHC staining results of the (B) C57/B6 and (D) BTBR representative sections. Images were centered on the sub-granular zone at 400x magnification.

The decreased blood flow in the ASD mouse model is consistent with the results seen in past research of the BTBR model [16]. Recent reviews have also shown supporting evidence for cerebral hypoperfusion from both PET/SPECT and fMRI studies [32]. The prevailing theory for this pathology is a dysfunctional vasodilation response in the autism cerebral arteries. A healthy brain will exhibit CBF that is largely unchanged over time due to the combined effect of the basal tone in smaller arterioles and the mechanical vascular resistance of larger arteries [33]. When the cerebral metabolic demands increase in an area, such as during an activity or task that recruits neurons from that brain region, the blood vessels dilate to reduce resistance and increase blood flow [34]. In ASD there appears to be a lack of this normal compensatory dilation and increase in blood flow when engaged in a task, such as during speaking or focusing on solving a problem [35,36]. When the blood vessels do not properly dilate, the smaller cross-sectional area of the lumen may cause higher blood pressure and restricted blood flow.

There also exist possible intrinsic structural explanations for these hypoperfusion results. Exacerbated inflammatory responses have been extensively documented in ASD patients. The resulting intracranial pressure can potentially cause mechanical vasoconstriction and as a result decreased blood flow [37,38]. Furthermore, evidence from recent research suggests that the majority of children with ASD also exhibit ongoing general neuroinflammatory disorders [39]. The increased absorption observed in the BTBR group could also be a result of neuro-inflammation. The increase in local tissue volume due to inflammation would result in increased chromophore concentration (in the form of hemoglobin and water) that would increase the absorption coefficient measurement. The lack of a significant difference in scattering parameter between the BTBR and C57/B6 models is contrary to expectations based on the decreased neurogenesis cell count shown in previous literature [16]. This could be due to a low sample size but could also be due to the physiological difference between the two mouse models. However, if we write µ_s_’ = *a*λ^−*b*^ for the Mie scattering representtaion, it is clear from the Figure 2B that b parameter (related to scattering power) is much smaller in autistic group (BTBR) than the control group (C57/B6). We will investigate this further by fitting both a and b from the measurements having additional wavelengths. Further investigation should be performed in terms of physiological measurements of the head anatomy between the two models.

## 4. Conclusion

The decreased CBF in the BTBR mouse model observed here is consistent with previous results. The depth-sensitive DCS device has shown that the blood flow in the autistic brain is significantly lower in the BTBR model. Frequency domain measurements showed differences in optical properties between the two models. Multimodal approach assessing tissue structure and function such as scattering coefficient, vascular structure, blood flow and blood volume measurements could give insights related to the assumption of dysfunctional vasodilation causing lower blood flow. We will pursue these studies in the near future.

## Funding

We thank the funding support from the Ohio Third Frontier (Sunar) to the Ohio Imaging Research and Innovation Network (OIRAIN, 667750).

## Disclosure

The authors declare that there are no conflicts of interest related to this article.

## Notes

### Competing Interest Statement

The authors have declared no competing interest.

## References

1. E. Simonoff, A. Pickles, T. Charman, S. Chandler, T. Loucas, and G. Baird, “Psychiatric disorders in children with autism spectrum disorders: Prevalence, comorbidity, and associated factors in a population-derived sample,” J. Am. Acad. Child Adolesc. Psychiatry 47(8), 921–929 (2008).

2. C. P. Johnson and S. M. Myers, “Identification and evaluation of children with autism spectrum disorders,” Pediatrics 120(5), 1183–1215 (2007).

3. C. Lord, S. Risi, L. Lambrecht, E. H. Cook, B. L. Leventhal, P. C. DiLavore, A. Pickles, and M. Rutter, “The Autism Diagnostic Observation Schedule-Generic: A standard measure of social and communication deficits associated with the spectrum of autism,” J. Autism Dev. Disord. 30(3), 205–223 (2000).

4. N. Boddaert and M. Zilbovicius, “Functional neuroimaging and childhood autism,” Pediatr. Radiol. 32(1), 1–7 (2002).

5. L. Galuska, S. Szakáll, M. Emri, R. Oláh, J. Varga, I. Garai, J. Kollár, I. Pataki, and L. Trón, “[PET and SPECT scans in autistic children],” Orv. Hetil. 143(21 Suppl 3), 1302–1304 (2002).

6. M. Sasaki, E. Nakagawa, K. Sugai, Y. Shimizu, A. Hattori, Y. Nonoda, and N. Sato, “Brain perfusion SPECT and EEG findings in children with autism spectrum disorders and medically intractable epilepsy,” Brain Dev. 32(9), 776–782 (2010).

7. M. Zilbovicius, B. Garreau, Y. Samson, P. Remy, C. Barthélémy, A. Syrota, and G. Lelord, “Delayed maturation of the frontal cortex in childhood autism,” Am. J. Psychiatry 152(2), 248–252 (1995).

8. M. Zilbovicius, N. Boddaert, P. Belin, J. B. Poline, P. Remy, J. F. Mangin, L. Thivard, C. Barthélémy, and Y. Samson, “Temporal lobe dysfunction in childhood autism: a PET study. Positron emission tomography,” Am. J. Psychiatry 157(12), 1988–1993 (2000).

9. G. S. Dichter, “Functional magnetic resonance imaging of autism spectrum disorders,” Dialogues Clin. Neurosci. 14(3), 319–351 (2012).

10. K. Jann, L. M. Hernandez, D. Beck-Pancer, R. McCarron, R. X. Smith, M. Dapretto, and D. J. J. Wang, “Altered resting perfusion and functional connectivity of default mode network in youth with autism spectrum disorder,” Brain Behav. 5(9), e00358 (2015).

11. R. C. M. Philip, M. R. Dauvermann, H. C. Whalley, K. Baynham, S. M. Lawrie, and A. C. Stanfield, “A systematic review and meta-analysis of the fMRI investigation of autism spectrum disorders,” Neurosci. Biobehav. Rev. 36(2), 901–942 (2012).

12. B. E. Yerys, J. D. Herrington, G. K. Bartley, H. S. Liu, J. A. Detre, and R. T. Schultz, “Arterial spin labeling provides a reliable neurobiological marker of autism spectrum disorder,” J. Neurodev. Disord. 10(1), (2018).

13. J. Li, L. Qiu, L. Xu, E. V Pedapati, C. A. Erickson, and U. Sunar, “Characterization of autism spectrum disorder with spontaneous hemodynamic activity,” Biomed. Opt. Express 7(10), 3871–3881 (2016).

14. J. Li, L. Qiu, L. Xu, E. V Pedapati, C. A. Erickson, and U. Sunar, “Characterization of autism spectrum disorder by functional near infrared spectroscopy (fNIRS),” in Biophotonics Congress: Biomedical Optics Congress 2018 (Microscopy/Translational/Brain/OTS), OSA Technical Digest (Optical Society of America, 2018), p. CTh2B.2.

15. J. Ouellette, X. Toussay, C. H. Comin, L. da F. Costa, M. Ho, M. Lacalle-Aurioles, M. Freitas-Andrade, Q. Y. Liu, S. Leclerc, Y. Pan, Z. Liu, J. F. Thibodeau, M. Yin, M. Carrier, C. J. Morse, P. Van Dyken, C. J. Bergin, S. Baillet, C. R. Kennedy, M. È. Tremblay, Y. D. Benoit, W. L. Stanford, D. Burger, D. J. Stewart, and B. Lacoste, “Vascular contributions to 16p11.2 deletion autism syndrome modeled in mice,” Nat. Neurosci. 23(9), (2020).

16. D. Abookasis, D. Lerman, H. Roth, M. Tfilin, and G. Turgeman, “Optically derived metabolic and hemodynamic parameters predict hippocampal neurogenesis in the BTBR mouse model of autism,” J. Biophotonics 11(3), (2018).

17. C. Poon, B. Rinehart, J. Li, and U. Sunar, “Cerebral Blood Flow-Based Resting State Functional Connectivity of the Human Brain using Optical Diffuse Correlation Spectroscopy,” JoVE (Journal Vis. Exp. (159), e60765 (2020).

18. K. K. Chadman, S. R. Guariglia, and J. H. Yoo, “New directions in the treatment of autism spectrum disorders from animal model research,” Expert Opin. Drug Discov. 7(5), 407–416 (2012).

19. D. W. Meechan, H. L. H. Rutz, M. S. Fralish, T. M. Maynard, L. A. Rothblat, and A.-S. LaMantia, “Cognitive Ability is Associated with Altered Medial Frontal Cortical Circuits in the LgDel Mouse Model of 22q11.2DS,” Cereb. Cortex (New York, NY) 25(5), 1143–1151 (2015).

20. D. T. Stephenson, S. M. O’Neill, S. Narayan, A. Tiwari, E. Arnold, H. D. Samaroo, F. Du, R. H. Ring, B. Campbell, M. Pletcher, V. A. Vaidya, and D. Morton, “Histopathologic characterization of the BTBR mouse model of autistic-like behavior reveals selective changes in neurodevelopmental proteins and adult hippocampal neurogenesis,” Mol. Autism 2(1), 7 (2011).

21. D. Wahlsten, P. Metten, and J. C. Crabbe, “Survey of 21 inbred mouse strains in two laboratories reveals that BTBR T/+ tf/tf has severely reduced hippocampal commissure and absent corpus callosum,” Brain Res. 971(1), 47–54 (2003).

22. C. Constantinides, R. Mean, and B. J. Janssen, “Effects of Isoflurane Anesthesia on the Cardiovascular Function of the C57BL/6 Mouse,” ILAR J. 52, e21–e31 (2011).

23. T. Durduran and A. G. Yodh, “Diffuse correlation spectroscopy for non-invasive, micro-vascular cerebral blood flow measurement,” Neuroimage 85 Pt 1, 51–63 (2014).

24. E. M. Buckley, A. B. Parthasarathy, P. E. Grant, A. G. Yodh, and M. A. Franceschini, “Diffuse correlation spectroscopy for measurement of cerebral blood flow: future prospects,” Neurophotonics 1(1), (2014).

25. Y. Lin, C. Huang, D. Irwin, L. He, Y. Shang, and G. Yu, “Three-dimensional flow contrast imaging of deep tissue using noncontact diffuse correlation tomography,” Appl. Phys. Lett. 104(12), 121103 (2014).

26. J. Li, C.-S. Poon, J. Kress, D. J. Rohrbach, and U. Sunar, “Resting-state functional connectivity measured by diffuse correlation spectroscopy,” J. Biophotonics 11(2), (2018).

27. R. C. Mesquita, T. Durduran, G. Yu, E. M. Buckley, M. N. Kim, C. Zhou, R. Choe, U. Sunar, and A. G. Yodh, “Direct measurement of tissue blood flow and metabolism with diffuse optics,” Philos. Trans. A. Math. Phys. Eng. Sci. 369(1955), 4390–4406 (2011).

28. G. Maret and P. E. Wolf, “Multiple light scattering from disordered media. The effect of brownian motion of scatterers,” Zeitschrift für Phys. B Condens. Matter 65(4), 409–413 (1987).

29. D. A. Boas, L. E. Campbell, and A. G. Yodh, “Scattering and imaging with diffusing temporal field correlations,” Phys. Rev. Lett. 75(9), 1855–1858 (1995).

30. D. A. Boas and A. G. Yodh, “Spatially varying dynamical properties of turbid media probed with diffusing temporal light correlation,” JOSA A 14(1), 192–215 (1997).

31. C. Cheung, J. P. Culver, K. Takahashi, J. H. Greenberg, and A. G. Yodh, “In vivo cerebrovascular measurement combining diffuse near-infrared absorption and correlation spectroscopies,” Phys. Med. Biol. 46(8), 2053–2065 (2001).

32. J. M. M. van der Burg, M. Björkqvist, and P. Brundin, “Beyond the brain: widespread pathology in Huntington’s disease,” Lancet. Neurol. 8(8), 765–774 (2009).

33. M. J. Cipolla, J. Smith, M. M. Kohlmeyer, and J. A. Godfrey, “SKCa and IKCa Channels, myogenic tone, and vasodilator responses in middle cerebral arteries and parenchymal arterioles: effect of ischemia and reperfusion,” Stroke 40(4), 1451–1457 (2009).

34. F. M. Faraci and D. D. Heistad, “Regulation of large cerebral arteries and cerebral microvascular pressure,” Circ. Res. 66(1), 8–17 (1990).

35. G. Allen and E. Courchesne, “Differential effects of developmental cerebellar abnormality on cognitive and motor functions in the cerebellum: an fMRI study of autism,” Am. J. Psychiatry 160(2), 262–273 (2003).

36. T. Ohnishi, H. Matsuda, T. Hashimoto, T. Kunihiro, M. Nishikawa, T. Uema, and M. Sasaki, “Abnormal regional cerebral blood flow in childhood autism,” Brain A J. Neurol. 123 (Pt 9, 1838–1844 (2000).

37. J. T. Morgan, G. Chana, C. A. Pardo, C. Achim, K. Semendeferi, J. Buckwalter, E. Courchesne, and I. P. Everall, “Microglial activation and increased microglial density observed in the dorsolateral prefrontal cortex in autism,” Biol. Psychiatry 68(4), 368–376 (2010).

38. D. L. Vargas, C. Nascimbene, C. Krishnan, A. W. Zimmerman, and C. A. Pardo, “Neuroglial activation and neuroinflammation in the brain of patients with autism,” Ann. Neurol. 57(1), 67–81 (2005).

39. J. K. Kern, D. A. Geier, L. K. Sykes, and M. R. Geier, “Relevance of Neuroinflammation and Encephalitis in Autism,” Front. Cell. Neurosci. 9, (2016).

